# Emergence of SARS-CoV-2 through Recombination and Strong Purifying Selection

**DOI:** 10.1101/2020.03.20.000885

**Authors:** Xiaojun Li, Elena E. Giorgi, Manukumar Honnayakanahalli Marichann, Brian Foley, Chuan Xiao, Xiang-Peng Kong, Yue Chen, Bette Korber, Feng Gao

**Affiliations:** Department of Medicine, Duke University Medical Center, Durham, NC 27710, USA; Theoretical Biology and Biophysics, Los Alamos National Laboratory, Los Alamos, NM 87544, USA; Department of Chemistry and Biochemistry, The University of Texas at El Paso, El Paso, TX 79968, USA; Department of Biochemistry and Molecular Pharmacology, Grossman School of Medicine, New York University, New York, NY 10016

## Abstract

COVID-19 has become a global pandemic caused by a novel coronavirus SARS-CoV-2. Understanding the origins of SARS-CoV-2 is critical for deterring future zoonosis and for drug discovery and vaccine development. We show evidence of strong purifying selection around the receptor binding motif (RBM) in the spike gene and in other genes among bat, pangolin and human coronaviruses, indicating similar strong evolutionary constraints in different host species. We also demonstrate that SARS-CoV-2’s entire RBM was introduced through recombination with coronaviruses from pangolins, possibly a critical step in the evolution of SARS-CoV-2’s ability to infect humans. Similar purifying selection in different host species and frequent recombination among coronaviruses suggest a common evolutionary mechanism that could lead to new emerging human coronaviruses.

**One Sentence Summary:** Extensive Recombination and Strong Purifying Selection among coronaviruses from different hosts facilitate the emergence of SARS-CoV-2

## Introduction

The severe respiratory disease COVID-19 was first noticed in late December 2019 (*1*). It rapidly became epidemic in China, devastating public health and finance. By mid-March, COVID-19 had spread to ∼150 countries and infected over 150,000 people (*2*). On March 11, 2020, the World Health Organization (WHO) officially declared it a pandemic.

A complete genome sequence of the etiological agent of COVID-19 (*3*), severe acute respiratory syndrome coronavirus 2 (SARS-CoV-2) (*4*), identified it as a new member of the genus *Betacoronavirus*, which include a diverse reservoir of coronaviruses (CoVs) isolated from bats (*5-7*). While genetically distinct from the betacoronaviruses that cause SARS and MERS in humans (*8, 9*), SARS-CoV-2 shares the highest level of genetic similarity (96.3%) with CoV RaTG13, sampled from a bat in Yunnan in 2013 (*8*). Recently, CoV sequences closely related to SARS-CoV-2 were obtained from confiscated Malaya pangolins in two separate studies (*10, 11*). Pangolin SARS-like CoVs (Pan_SL-CoV) form two distinct clades corresponding to their collection location. Pan_SL-CoV_GD from Guangdong (GD) province in China and are genetically more similar to SARS-CoV-2 (91.2%) than Pan_SL-CoV_GX from Guangxi (GX) province (85.4%).

Understanding the origin of SARS-CoV-2 may help resolve strategies to deter future cross-species transmissions and to establish appropriate animal models. Viral sequences nearly identical to SARS and MERS viruses were found in civets and domestic camels, respectively (*12, 13*), demonstrating that they originated from zoonotic transmissions with intermediate host species between the bat reservoirs and humans—a common pattern leading to CoV zoonosis (*5*). Viruses nearly identical to SARS-CoV-2 have not yet been found. In this paper we demonstrate, through localized genomic analysis, a complex pattern of evolutionary recombination between CoVs from distinct host species and cross-species infections that likely originated SARS-CoV-2.

## Results

### Acquisition of receptor binding motif through recombination

Phylogenetic analysis of 43 complete genome sequences from three clades (SARS-CoVs and bat_SL-CoVs; SARS-CoV-2, bat_SL-CoVs and pan_SL-CoVs; and two divergent bat_SL-CoVs) within the Sarbecovirus group (*9*) confirms that RaTG13 is overall the closest sequence to SARS-CoV-2 (fig. S1). It is followed by Pan_SL-CoV_GD viruses next, and then Pan_SL-CoV_GX. Among the bat-CoV sequences in clade 2 (fig. S1), ZXC21 and ZC45, sampled from bats in 2005 in Zhoushan, Zhejiang, China, are the most divergent, with the exception of the beginning of the ORF1a gene (region 1, fig. 1A). All other Bat_SL-CoV and SARS-CoV sequences form a separate clade 3, while clade 1 comprises BtKY72 and BM48-31, the two most divergent Bat_SL-CoV sequences, in the Sarbecovirus group (fig. S1). Recombination in the first SARS-CoV-2 sequence (Wuhan-Hu-1) with other divergent CoVs has been previously observed (*3*). Here, to better understand the role of recombination in the origin of SARS-CoV-2 among these genetically similar CoVs, we compare Wuhan-Hu-1 to six representative Bat_SL-CoVs, one SARS-CoV, and the two Pan_SL-CoV_GD sequences using SimPlot analysis (*14*). RaTG13 has the highest similarity across the genome (*8*), with two notable exceptions where a switch occurs (fig. 1A). In phylogenetic reconstructions, SARS-CoV-2 clusters closer to ZXC21 and ZC45 than RaTG13 at the beginning of the ORF1a gene (region 1, fig. 1B), and, as reported (*10, 15*), to a Pan_SL-CoV_GD in region 2 (fig.s 1C and S2), which spans the receptor angiotensin-converting enzyme 2 (ACE2) binding site in the spike (S) glycoprotein gene. Comparing Wuhan-Hu-1 to Pan_SL-CoV_GD and RaTG13, as representative of distinct host-species branches in the evolutionary history of SARS-CoV-2, using the recombination detection tool RIP (*16*), we find significant recombination breakpoints before and after the ACE2 binding site (fig. S2A), suggesting that SARS-CoV-2 carries a history of cross-species recombination between the bat and the pangolin CoVs.

**Fig. 1.**
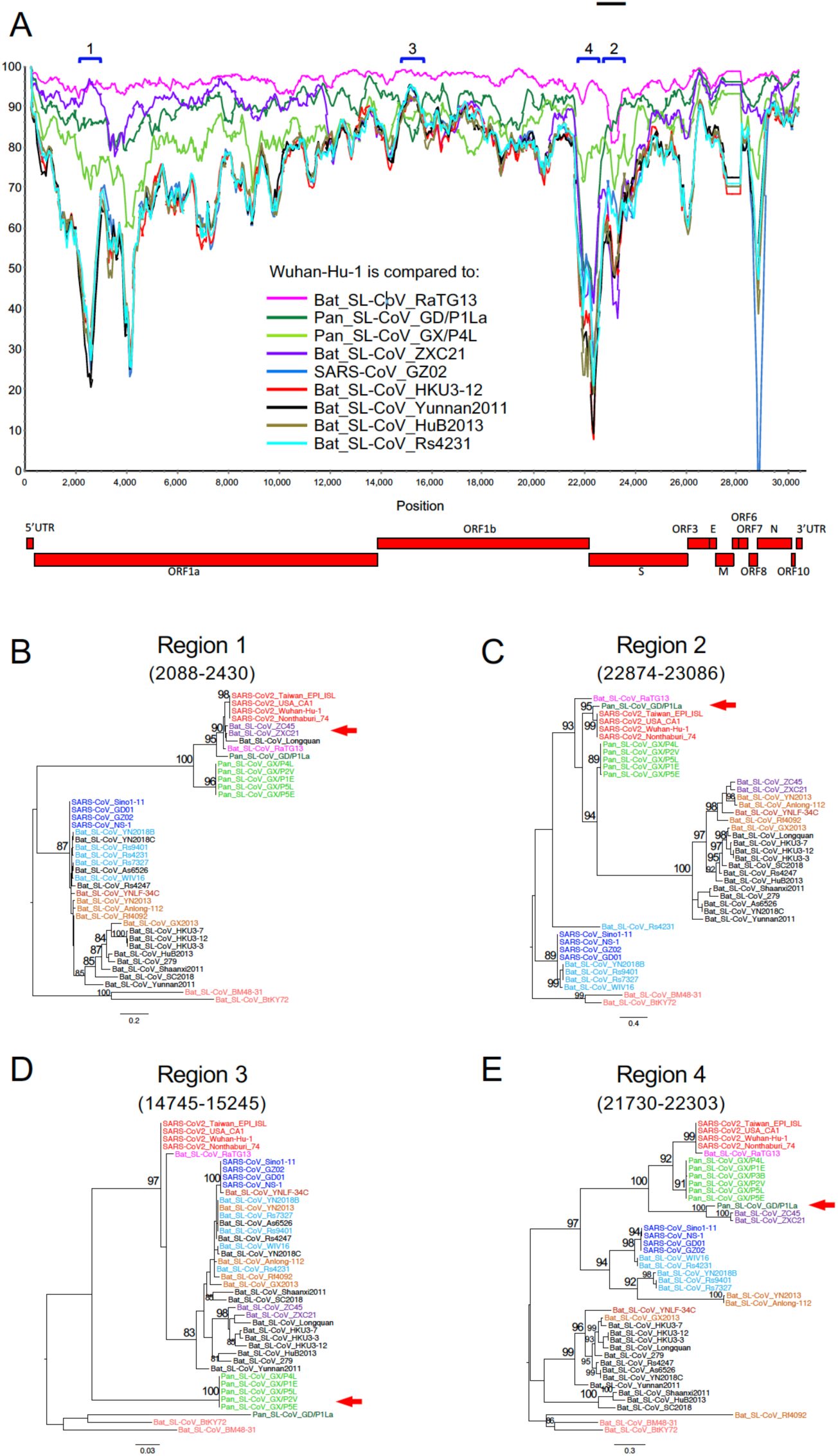
SARS-CoV-2 recombination with Pan_SL-CoV and Bat_SL-CoV. (A) SimPlot genetic similarity plot between SARS-CoV-2 Wuhan-Hu-1 and representative CoV sequences, using a 400-bp window at a 50-bp step and the Kimura 2-parameter model. (B-E) Phylogenetic trees of regions of disproportional similarities, showing high similarities with ZXC21 (B) or GD/P1La (C), or high divergences with both GD/P1La and GX/P4L (D) or GD/P1La (E). All positions are relative to Wuhan-Hu-1.

Pan_SL-CoV sequences are generally more similar to SARS-CoV-2 than CoV sequences, other than RaTG13 and ZXC21, but are more divergent from SARS-CoV-2 at two regions in particular: the beginning of the ORF1b gene and the highly divergent N terminus of the S gene (regions 3 and 4, respectively, fig. 1A). Within-region phylogenetic reconstructions show that Pan_SL-CoV sequences become as divergent as BtKY72 and BM48-31 in region 3 (fig. 1D), while less divergent in region 4, where Pan_SL-CoV_GD clusters with ZXC21 and ZC45 (fig. 1E). Together, these observations suggest ancestral cross-species recombination between pangolin and bat CoVs in the evolution of SARS-CoV-2 at the ORF1a and S genes. Furthermore the discordant phylogenetic clustering at various regions of the genome among clade 2 CoVs also supports extensive recombination among these viruses isolated from bats and pangolins.

The SARS-CoV-2 S glycoprotein mediates viral entry into host cells and therefore represents a prime target for drug and vaccine development (*17, 18*). While SARS-CoV-2 sequences share the greatest overall genetic similarity with RaTG13, this is no longer the case in parts of the S gene. Specifically, amino acid sequences of the receptor binding motif (RBM) in the C terminal of the S1 subunit are nearly identical to those in two Pan_SL-CoV_GD viruses, with only one amino acid difference (Q498H)—although the RBM region has not been fully sequenced in one of Guangdong pangolin virus (Pan_SL-CoV_GD/P2S) (fig. 2A). Pangolin CoVs from Guangxi are much more divergent. Phylogenetic analysis based on the amino acid sequences of this region shows three distinct clusters of SARS-CoV, SARS-CoV-2 and bat-CoV only viruses, respectively (fig. 2B). Interestingly, while SARS-CoV and SARS-CoV-2 viruses use ACE2 for viral entry, all CoVs in the third cluster have a 5-aa deletion and a 13-14-aa deletion in RBM (fig. 2A) and cannot infect human target cells (*5, 19*).

**Fig. 2.**
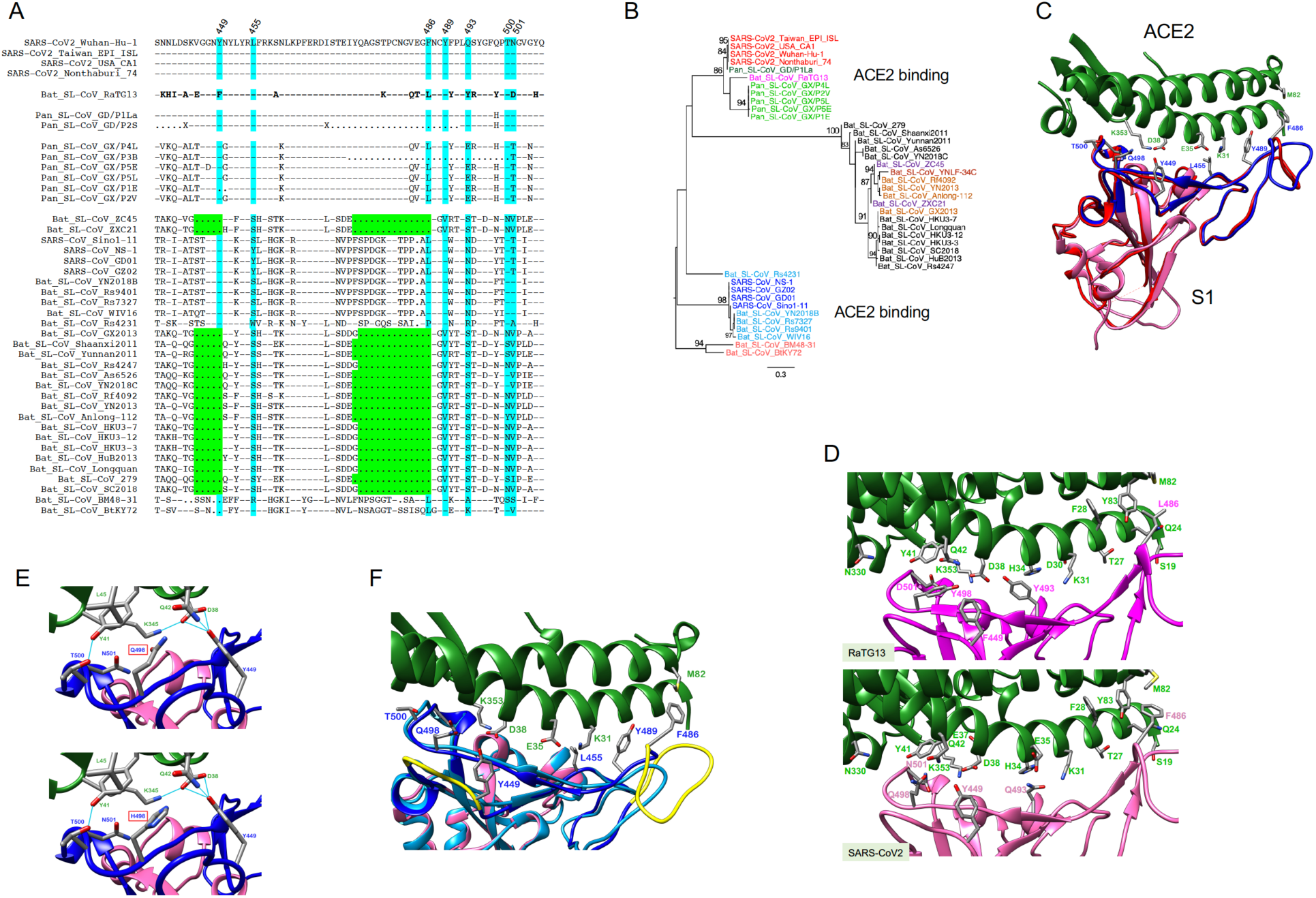
Impact of SARS-CoV-2 recombination on coreceptor binding. (A) AA sequences of the receptor binding motif (RBM) in the spike (S) gene among Sarbecovirus CoVs compared to Wuhan-Hu-1 (top). Dashes indicate identical aa’s, dots indicate deletions. ACE2 critical contact sites highlighted in blue, two large deletions in green. (B) RBM aa phylogenetic tree, showing three distinct clusters, with large deletions Bat-SL-CoVs in divergent cluster. (C) SARS-CoV and SARS-CoV-2 receptor binding domains (RBD). Human ACE2 in green at the top and the S1 unit of the S-protein at the bottom; SARS-CoV S-protein (PDB 2AJF) in red, and SARS-CoV-2 S-protein in magenta with RBM in blue. All structure backbones shown as ribbons with key residues at the interface shown as stick models, labeled using the same color scheme. (D) Impact of different RBM aa between SARS-CoV-2 RaTG13 on ACE2 binding. (E) Impact of different aa at position 498 (Q in SARS-CoV-2, top, and H in RaTG13, bottom) on ACE2 binding. Same color-coding as in (C) with additional hydrogen bonding as light blue lines. (F) Impact of two deletions on ACE2 binding interface in some bat-SL-CoVs, positions indicated in yellow, and modeled structure with long deletion between residue 473 in light blue.

Although both SARS-CoV and SARS-CoV-2 use the human ACE2 as their receptors (*8, 20*) they show a high level of genetic divergence (figs. 1 and S1). However, structures of the S1 unit of the S protein from both viruses are highly similar (*21-23*), with the exception of a loop, not proximal to the binding site, that bends differently (fig. 2C). This suggests that viral entry through binding of ACE2 is structurally constrained to maintain the correct conformation. Among 17 distinct amino acids between SARS-CoV-2 and RaTG13 (fig. 2A), five contact sites are different, likely impacting RaTG13’s binding to ACE2 (fig. 2D and Table S1). The single amino acid difference (Q or H at position 498) between SARS-CoV-2 and Pan_SL-CoV_GD is at the edge of the ACE2 contact interface; neither Q or H at this position form hydrogen bonds with ACE2 residues (fig. 2E). Thus, a functional RBM nearly identical to the one in SARS-CoV-2 is naturally present in Pan_SL-CoV_GD viruses. The very distinctive RaTG13 RBM suggests that this virus is unlikely to infect human cells, and that the acquisition of a complete functional RBM by a RaTG13-like CoV through a recombination event with a Pan_SL-CoV_GD-like virus enabled it to use ACE2 for human infection.

Three small insertions are identical in SARS-CoV-2 and RaTG13 but not found in other CoVs in the Sarbecovirus group (*24*). The RaTG13 sequence was sampled in 2013, years before SARS-CoV-2 was first identified. It is unlikely that both SARS-CoV-2 and RaTG13 independently acquired identical insertions at three different locations in the S gene. Thus, it is plausible that an RaTG13-like virus served as a progenitor to generate SARS-CoV-2 by gaining a complete human ACE2 binding RBM from Pan_SL-CoV_GD-like viruses through recombination. Genetic divergence at the nucleic acid level between Wuhan-Hu-1 and Pan_SL-CoV_GD viruses is significantly reduced from 13.9% (fig. 1E) to 1.4% at the amino acid level (fig. 2B) in the RBM region, indicating recombination between RaTG13-like CoVs and Pan_SL-CoV_GD-like CoVs. Furthermore, SARS-CoV-2 has a unique furin cleavage site insertion (PRRA) not found in any other CoVs in the Sarbecovirus group (*24*), although similar motifs are also found in MERS and more divergent bat CoVs (*25*) (Fig. S3). This PRRA motif makes the S1/S2 cleavage in SARS-CoV-2 much more efficiently than in SARS-CoV and may expand its tropism and/or enhance its transmissibility (*23*). A recent study of bat CoVs in Yunnan, China, identified a three-amino acid insertion (PAA) at the same site (*26*). Although it is not known if this PAA motif can function like the PRRA motif, the presence of a similar insertion at the same site indicates that such insertion may already be present in the wild bat CoVs. The more efficient cleavage of S1 and S2 units of the spike glycoprotein (*25*) and efficient binding to ACE2 by SARS-CoV-2 (*22, 27*) may have allowed SARS-CoV-2 to jump to humans, leading to the rapid spread of SARS-CoV-2 in China and the rest of the world.

### Strong purifying selection among SRAS-CoV-2 and closely related viruses

Recombination from Pan_SL-CoV_GD at the RBM and at the unique furin cleavage site insertion prompted us to examine the SARS-CoV-2 sequences within these regions. Amino acid sequences from SARS-CoV-2, RaTG13, and all Pan_SL-CoV viruses are identical or nearly identical before, between, and after the RBM and the furin cleavage site, while all other CoVs are very distinctive (fig. 3A and S3). The average of all pairwise dN/dS ratios (ω) among SARS-CoV-2, RaTG13, and Pan_SL-CoV viruses at the 3’-end of the S gene (after the furin cleavage site) is 0.013, compared to ω =0.05 in the S region preceding the furin cleavage site, and to ω =0.04 after the site for all other CoVs. The much lower ω value at the 3’-end of the S gene among the SARS-CoV-2, RaTG13, and Pan_SL-CoV viruses indicates that this region is under strong purifying selection within these sequences (fig. 3A). A plot of synonymous and nonsynonymous substitutions relative to Wuhan-Hu-1 highlights the regional differences across the region before and after the RBM and the furin cleavage site (fig. 3A): the 3’ end of the region is highly conserved among the SARS-CoV-2, RaTG13, and Pan_SL-CoV viruses (Group A), while far more nonsynonymous mutations are observed in the rest of the CoV sequences (Group B). The shift in selective pressure in the 3’ -end of the gene among these related viruses versus other CoVs begins near codon 368 (fig. 3B), and such a shift was not evident among other compared CoVs (fig. 3B-D).

**Fig. 3.**
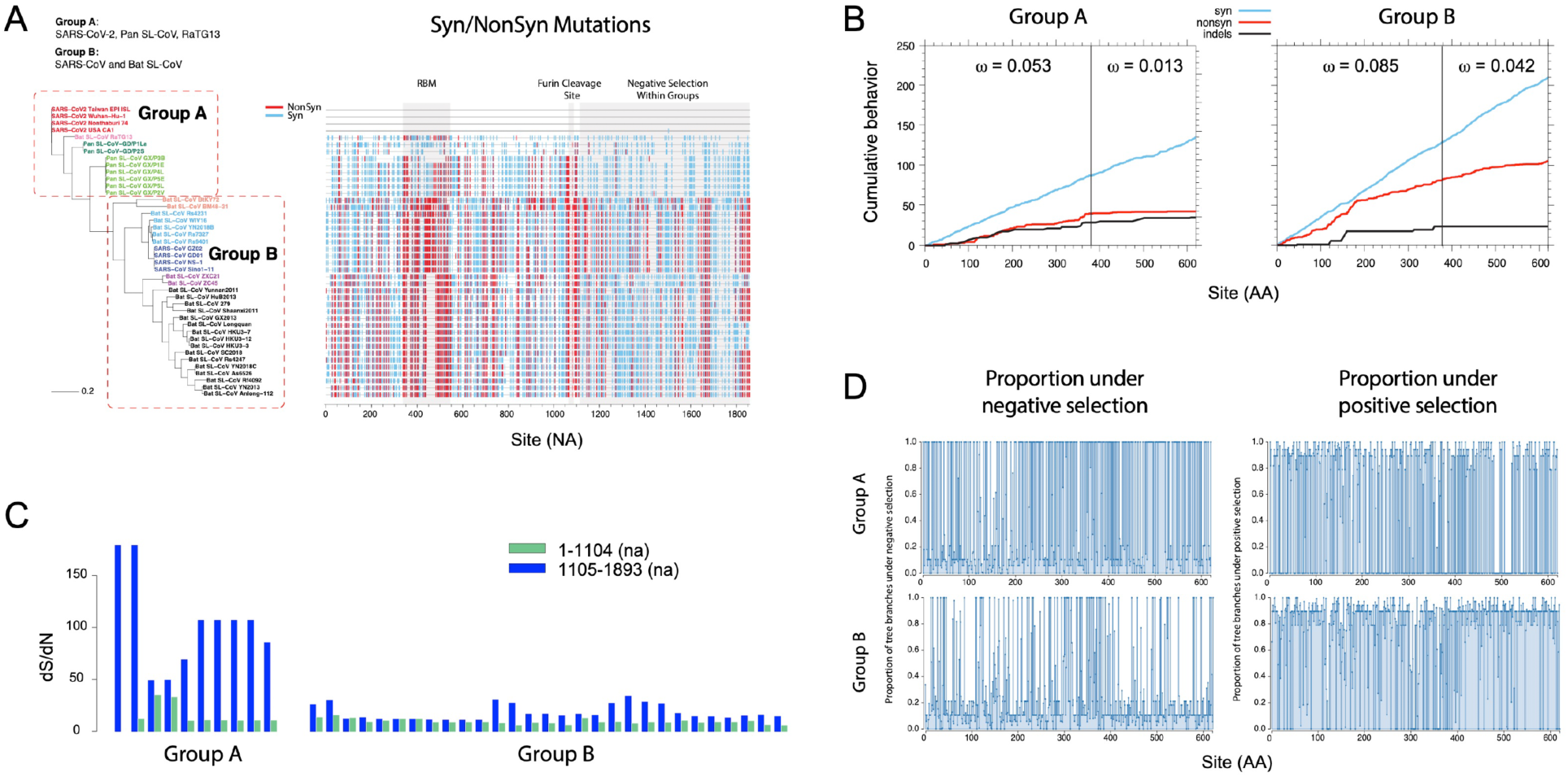
Strong purifying selection after furin cleavage in S gene among SARS-CoV-2 and closely related viruses. (A) Phylogenetic tree (left) and Highlighter plot (right) of sequences around the RBM and furin cleavage site compared to SARS-CoV-2 Wuhan-Hu-1 (na positions 22541-24391). ACE2 receptor binding motif (RBM) and furin cleavage site highlighted in light-gray boxes. Mutations compared to Wuahn-Hu-1 are light blue for synonymous, red for non-synonymous. Dominance of synonymous mutations within group A compared to group B highlighted on the right. (B) Cumulative plots of each codon average behavior for all pairwise comparisons for indels and synonymous (light blue) and non-synonymous (red) mutations, by group. A vertical steps in group A at around codon 370 (na 1105) shows a shift in localized accumulations; non-synonymous mutations end after the furin cleavage site. Group B instead lacks this abrupt change in slope. ω’s denote average ratios of the rate of nonsynonymous substitutions per nonsynonymous site (dN/dS) for each group and region half. (C) Sequence dS/dN ratios compared to Wuhan-Hu-1 within codons 1-370 (na 1-1104, green) and codons 371-620 (na 1105-1893, dark blue). (D) Proportion of tree branches under positive and negative selection (right and left respectively) per site for the two groups using the mixed effects model of evolution (MEME) from datamonkey (www.datamonkey.org).

We observe similar patterns of purifying selection pressure in other parts of the genome, including the E and M genes, as well as the partial ORF1a and ORF1b genes (fig. 4). Interestingly, the purifying selection pressure varies among different viruses depending on which genes are analyzed. The broadest group includes SARS-CoV-2, RaTG13, all Pan_SL-CoV and the two bat CoVs (ZXC21 and ZC45) for both E and M genes (figs. 4 and S5). The second group includes SARS-CoV-2, RaTG13, and all Pan_SL-CoV only for the 3’ end of the S gene. The narrowest selection group only contains SARS-CoV-2, RaTG13, and pangolin CoVs from Guangdong for the partial regions of ORF1a and ORF1b (figs. 4 and S6). Consistently low ω values and strong purifying selection pressure on SARS-CoV-2, RaTG13 and Pan_SL-CoV_GD viruses suggest that these complete and partial genes are under similar functional/structural constraints among the different host species. In two extreme cases, amino acid sequences of the E gene and the 3’ end of ORF1a are identical among the compared CoV sequences, although genetic distances are quite large among these viruses at the nucleic acid level. Such evolutionary constraints across viral genomes, especially at functional domains in the S gene, which plays an important role in cross-species transmission (*5, 17*), coupled with frequent recombination, may facilitate cross-species transmissions between RaTG13-like bat and/or Pan_SL-CoV_GD-like viruses.

**Fig. 4.**
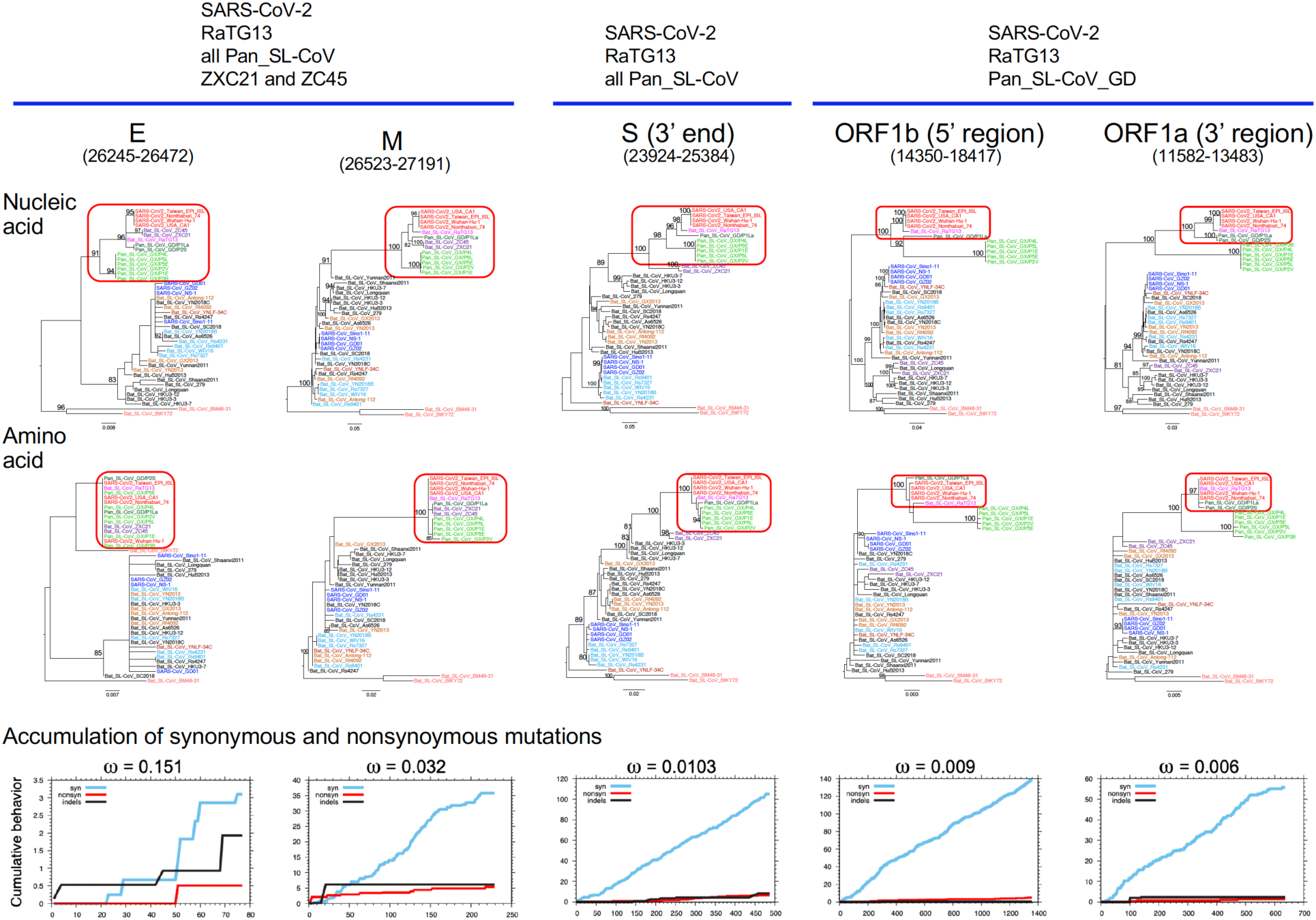
Strong purifying selection on complete and partial gene regions among SARS-CoV-2, RaTG13 and Pan_SL-CoV viruses. Purifying selection pressure on complete and partial genes within different viruses (red boxes) as evident by shorter branches in aa phylogenetic trees compared to na trees. Distinct purifying selection patterns are observed among different viruses: (A) SARS-CoV-2, RaTG13, Pan_SL-CoV and bat CoV ZXC21 and ZC45; (B) SARS-CoV-2, RaTG13, all Pan_SL-CoV sequences; (C) SARS-CoV-2, RaTG13 and Pan_SL-CoV_GD. Cumulative plots of the average behavior of each codon for all pairwise comparisons for synonymous mutations, non-synonymous mutations and indels within each gene region. ω’s denote average ratios of the rate of nonsynonymous substitutions per nonsynonymous site (dN/dS) for each group.

### Frequent recombination between SARS-CoVs and bat_SL-CoVs

Previous studies using more limited sequence sets found that SARS-CoVs originated through multiple recombination events between different bat-CoVs (*10, 17, 19, 28, 29*). Our phylogenetic analyses of individual genes show that SARS-CoV sequences tend to cluster with YN2018B, Rs9401, Rs7327, WIV16 and Rs4231 (group A) for some genes and Rf4092, YN2013, Anlong-112 and GX2013 (group B) for others (fig. S7). SimPlot analysis using both groups of bat_SL-CoVs and the closely related bat CoV YNLF-34C (*29*) shows that SARS-CoV GZ02 shifts in similarity across different bat SL-CoVs at various regions of the genome (fig. 5A). In particular, phylogenetic reconstruction of the beginning of ORF1a (region 1) confirms that SARS-CoVs cluster with YNLF-34C (*29*), and this region is distinctive comparing to all other CoVs (fig. 5B). YNLF-34C is more divergent from SARS-CoV than other bat-CoV viruses before and after this region, confirming the previously reported complex recombinant nature of YNLF-34C (*29*) (fig. 5A). At the end of the S gene (region 2), SARS-CoVs cluster with group A CoVs, forming a highly divergent clade (fig. 5C). In region 3 (ORF8), SARS-CoVs and group B CoVs, together with YNLF-34C, form a very divergent and distinctive cluster (fig. 5D). To further explore the recombinant nature of SARS-CoVs, we compared GZ02 to representative bat CoV sequences using the RIP recombination detection tool (*16*). We identified four significant breakpoints (at 99% confidence) between the two parental lineages (fig. S8A), further supported by phylogenetic analysis (fig. S8B-S8D). In addition, the two aforementioned groups of bat CoVs (shown in light brown and light blue in the trees) show similar cluster changes across the five recombinant regions, suggesting multiple events of historic recombination among bat SL-CoVs. These results demonstrate that SARS-CoV shares a recombinant history with at least three different groups of bat-CoVs and confirms the major role of recombination in the evolution of these viruses.

**Fig. 5.**
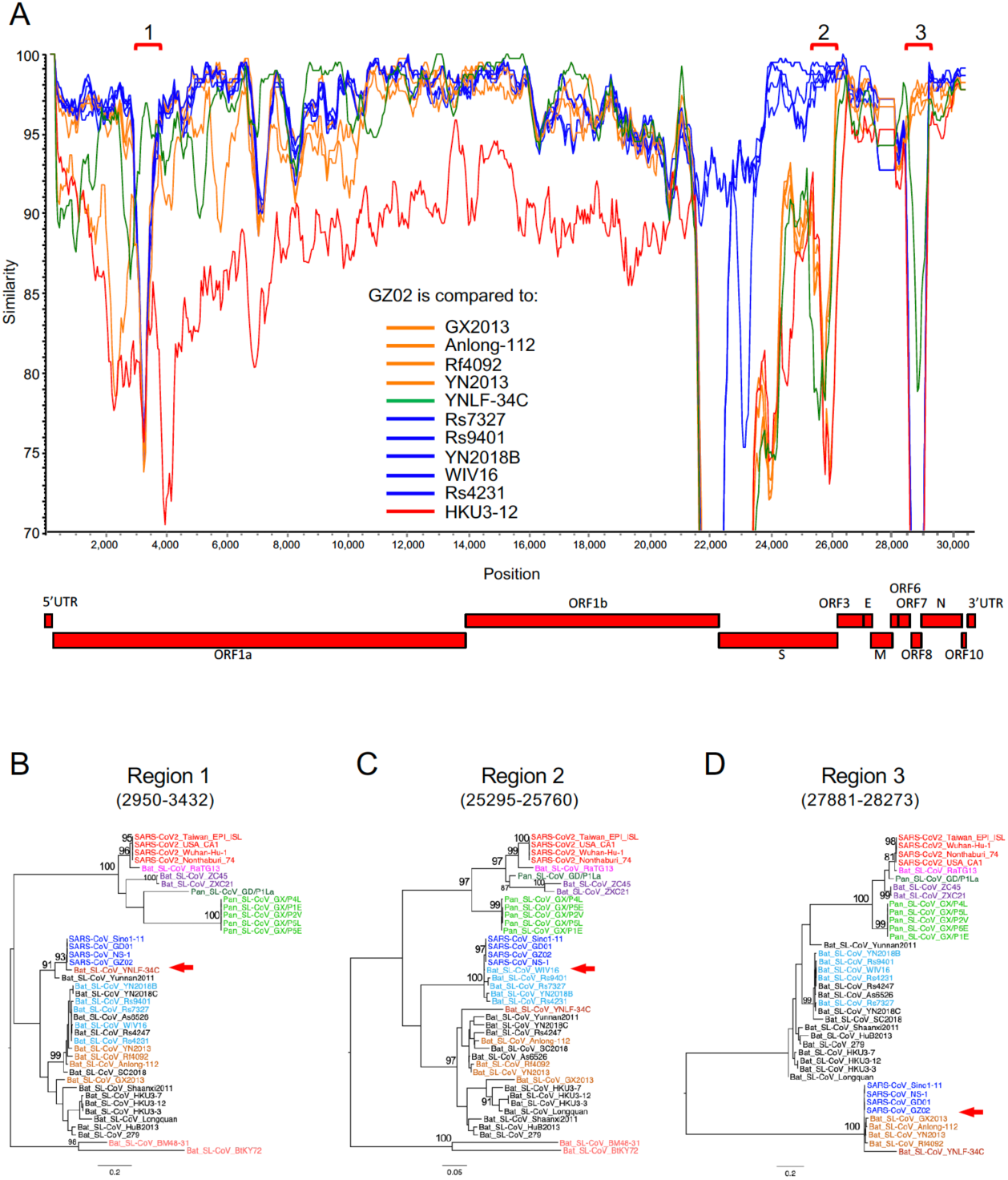
Multiple recombination of SARS-CoVs with different bat_SL-CoVs. (A) SimPlot genetic similarity plot between SARS-CoV GZ02 and SARS_SL-CoVs, using a 400-bp window at a 50-bp step and the Kimura 2-parameter model. Group A reference CoVs (YN2018B, Rs9401, Rs7327, WIV16 and Rs4231) in blue, group B CoVs (Rf4092, YN2013, Anlong-112 and GX2013) in orange, YNLF-34C in green, and outlier control HKU3-12 in red. (B-D) Phylogenetic trees for high similarity regions between GZ02 and YNLF-34C (B), group (C), and group B (D). All positions are relative to Wuhan-Hu-1.

Of the bat SL-CoVs that contributed to the recombinant origin of SARS-CoV, only group A viruses bind to ACE2. Group B bat SL-CoVs do not infect human cells (*5, 19*) and have two deletions in the RBM (figs. 1E and 2A). The short deletion between residues 445 and 449, and in particular the loss of Y449, which forms three hydrogen bonds with ACE2, will significantly affect the overall structure of the RBM (figs. 2F and 2E). The region encompassing the large deletion between residues 473 and 486 contains the loop structure that accounts for the major differences between the S protein of SARS-CoV and SARS-CoV-2 (fig. 2C). This deletion causes the loss of contact site F486 and affects the conserved residue F498’s hydrophobic interaction with residue M82 on ACE2 (fig. 2F). These two deletions will render RBM in those CoVs incapable to bind human ACE2. Therefore, recombination may play a role in enabling cross-species transmission in SARS-CoVs through the acquisition of an S gene type that can efficiently bind to the human ACE2 receptor.

ORF8 is one of the highly variable genes in coronaviruses (*5, 17, 29*) and its function has not yet been elucidated (*5, 17, 30*). Breakpoints within this region show that recombination occurred at the beginning and the end of ORF8 (fig. S9), where nucleic acid sequences are nearly identical among both SARS-CoVs and group B bat CoVs. Moreover, all compared viruses form three highly distinct clusters (fig. 5D), suggesting that the ORF8 gene may be biologically constrained and evolves through modular recombination. The third recombination region at the beginning of ORF1a is where SARS-CoV-2 also recombined with other bat CoVs (region 1, fig. 1A). This region is highly variable (*5, 17*) and recombination within this part of the genome was also found in many other CoVs, suggesting that it may be a recombination hotspot and may factor into cross-species transmission.

## Discussion

There are three important aspects to betacoronavirus evolution that should be carefully considered in phylogenetic reconstructions among more distant coronaviruses. First, there is extensive recombination among all of these viruses (*10, 17, 19, 28, 29*) (figs. 1 and 5), making standard phylogenetic reconstructions based on full genomes problematic, as different regions of the genome have distinct ancestral relationships. Second, between more distant sequences, synonymous substitutions are often fully saturated, which can confound analyses of selective pressure and add noise to phylogenetic analysis. Finally, there are different selective pressures at work in different lineages, which is worth consideration interpreting trees.

The currently sampled pangolin CoVs are too divergent from SARS-CoV-2 for them to be SARS-CoV-2 progenitors, but it is noteworthy that these sequences contain an RBM that can most likely bind to human ACE2. While RaTG13 is the most closely related CoV sequence to SARS-CoV-2, it has a distinctive RBM, which is not expected to bind to human ACE2. SARS-CoV-2 has a nearly identical RBM to the one found in the pangolin CoVs from Guangdong. Thus, it is plausible that RaTG13-like bat-CoV viruses may have obtained the RBM sequence binding to human ACE2 through recombination with Pan_SL-CoV_GD-like viruses. We hypothesize that this, and/or other ancestral recombination events between viruses infecting bats and pangolins, may have had a key role in the evolution of the strain that lead to the introduction of SARS-CoV-2 into humans.

All three human CoVs (SARS, MERS and SARS-2) are the result of recombination among CoVs. Recombination in all three viruses involved the S gene, likely a precondition to zoonosis that enabled efficient binding to human receptors (*5, 17*). Extensive recombination among bat coronaviruses and strong purifying selection pressure among viruses from humans, bats and pangolin may allow such closely related viruses ready jump between species and adapt to the new hosts. Many bat CoVs have been found able to bind to human ACE2 and replication in human cells (*10, 19, 31-33*). Serological evidence has revealed that additional otherwise undetected spillovers have occurred in people in China living in proximity to wild bat populations (*34*). Continuous surveillance of coronaviruses in their natural hosts and in humans will be key to rapid control of new coronavirus outbreaks.

So far efforts have failed to find the original pathway of SARS-CoV-2 into humans by identifying a coronavirus that is nearly identical to SARS-CoV-2, as those found for SARS and MERS in civets and domestic camels respectively (*12, 13*). However, if the new SARS-CoV-2 strain did not cause widespread infections in its natural or intermediate hosts, such a strain may never be identified. The close proximity of animals of different species in a wet market setting may increase the potential for cross-species spillover infections, by enabling recombination between more distant coronaviruses and the emergence of recombinants with novel phenotypes. While the direct reservoir of SARS-CoV-2 is still being sought, one thing is clear: reducing or eliminating direct human contact with wild animals is critical to preventing new coronavirus zoonosis in the future.

## Materials and Methods

### Sequences analysis

All 43 CoV complete genome sequences were obtained from GenBank and GISAID (Global Initiative on Sharing All Influenza Data) (*35, 36*), and were selected to be representative of the diversity. Pan_SL-CoV_GD/P1La sequence was generated by combining Pan_SL-CoV_GD/P1L (*10*) with some additional sequences from the NCBI BioProject database PRJNA5732983 (*11, 37*) to have a maximal coverage of the complete genome sequence for analysis. A new CoV sequence from pangolin (EPI_ISL_410721) (*38*)was not inclued because that it became available after we had already completed the analyses in this study, and it was not as close to SARS-CoV-2 sequences and did not change the interpretation of our results. The whole genome sequences were first aligned using Clustal X2 (*39*). The alignments for all coding regions were manually optimized based on the amino acid sequence alignment using SeaView 5.0.1.

### Recombination Analyses

SimPlot 3.5.15 (*14*) was used determine the percent identity of the query sequence to reference sequences. Potential recombinant regions among analyzed sequences were identified by sliding a 400bp-window at a 50bp-step across the alignment using the Kimura 2-parameter model. Phylogenetic trees were constructed by the maximum likelihood method using the GTR model (*40*), and their reliability was estimated from 1,000 bootstrap replicates. The positions of analyzed sequence regions were based on those in the reference SARS-CoV-2 Wuhan-Hu-1 (MN908947). Recombination regions and breakpoints were also analyzed using the LANL database (*41*) tool RIP (*16*).

### Selection Analyses

Cumulative plots of the average behavior of each codon for all pairwise comparisons in the input data, for insertions and deletions (indels), synonymous (syn), and nonsynonymous (nonsyn) mutations and values of the ratios of the rate of synonymous nucleotide substitutions per synonymous site and nonsynonymous substitutions per nonsynonymous site (dN/dS, or ω) were obtained using the LANL database tool SNAP (*42*). In order to avoid counting instances where synonymous mutations were saturated, averages of all pairwise dN/dS ratios were calculated excluding pairs that yielded dS values greater than 1. Sequences were analyzed for episodic selection pressure using the mixed effects model of evolution (MEME) (*43*) from the datamonkey server (www.datamonkey.org).

### Structure modeling of receptor binding

To investigate the single mutation Q498H in RBM between SARS-CoV-2 and Pan_SL-CoV_GD, Q498 in the crystal structure of S/ACE2 complex was mutated to H498 using Chimera (*44*). Local energy minimization (only H498 was allowed to move) was computed using Chimera’s built-in functions. To investigate the impact of the deletion between residue 473 to 486 to the binding interface between SARS-CoV-2 and human ACE2, a homology model with the deletion was generated using I-TASSER (*45*). The top five best models provided by the server have Confidence Score (C-score) of 0.86, -2.33, -4.01, -4.17, and -4.49. The C-score was used to estimate the quality of the models, which should be between -5.0 to 2; the higher the value, the higher the confidence in the model (*45*). Based on the C-score, model 1 was used in Figure 2F. The interaction of the RBD of RaTG13 and ACE2 was modeled on PDB 6VW1, a hybrid structure of human SARS-CoV2 (*46*) using ICM software package (*35*), and the mutational differences of the Gibbs free energy (Table S1) were calculated with the built-in algorithm.

## Acknowledgments

We thank all those who have contributed SARS-CoV-2 genome sequences to the GISAID database (https://www.gisaid.org/) and Virological.org (http://virological.org/). We also thank Dr. Xinquan Wang from Tsinghua University for sharing the PDB 6M0J structure with us before its official release date. Funding: This work was supported by NIH Grants (AI122909, AI118571, GM129525 and AI145655). EEG, BK, and BF acknowledge support by the LDRD program at Los Alamos National Laboratory. Author contributions: Project conceptualization: F.G., B.K., E.E.G; Structure analysis: C.X., X-P.K; Sequence analysis: F.G., B.K., X.L., E.E.G., M.H.M., Y.C., B.F; Phylogenetic analysis: F.G., B.K., X.L., E.E.G., M.H.M., Y.C.; Recombination analysis: F.G., E.E.G., B.K., X.L., E.E.G., M.H.M., B.F.; Manuscript writing: F.G., B.K., E.E.G. Manuscript editing: F.G., B.K., E.E.G., X.L., C.X., X-P.K., F.G. and B.F. supervised the project. Competing interests: All authors declare no competing interests. All data is available in the main text or the supplementary materials.

## Supplementary Material

Materials and Methods

Figures S1-S9

Table S1

References (35-46)

## Notes

#### Summary of Updates

author contributions and methods updated

