## Supplemental files for "Emergence of SARS-CoV-2 through Recombination and Strong Purifying Selection"

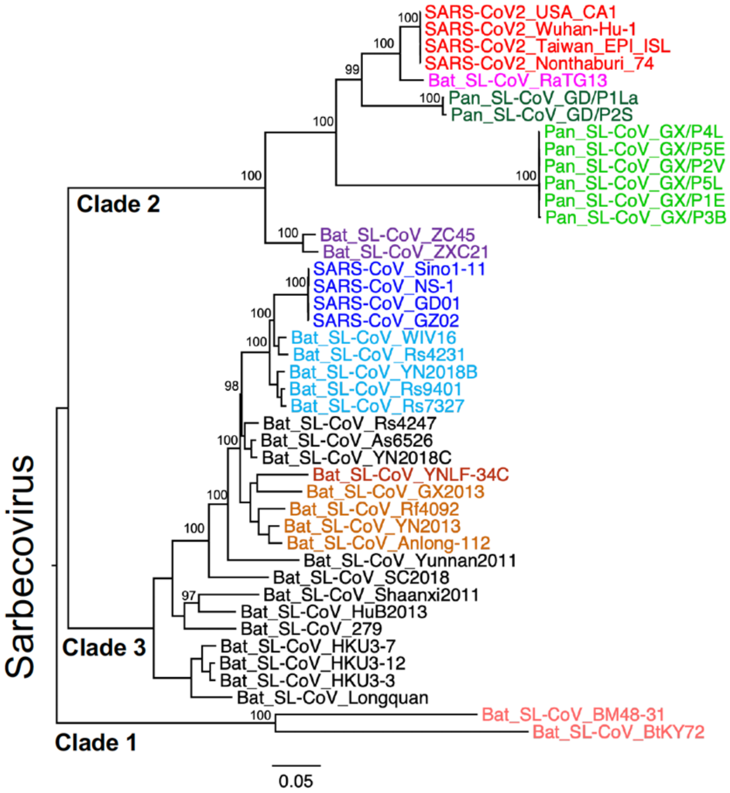


**Fig. S1. Phylogenetic tree of the complete CoV genome sequences.** All 43 sequences used in this study includes: 4 SARS-CoV-2 sequences (red), Bat_SL-CoV sequence RaTG13 (magenta), 2 pangolin CoV from Guangdong (Pan_SL-CoV_GD, dark green), 6 pangolin CoV from Guangxi (Pan_SL-CoV_GX , light green), and 4 SARS-CoV sequences (dark blue). The remaining Bat_SL-CoV sequences in the set are color-coded according to their phylogenetic subclusterings in the tree. Phylogenetic trees were constructed by the maximum likelihood method using the GTR model (*8*), and their reliability was estimated from 1,000 bootstrap replicates.

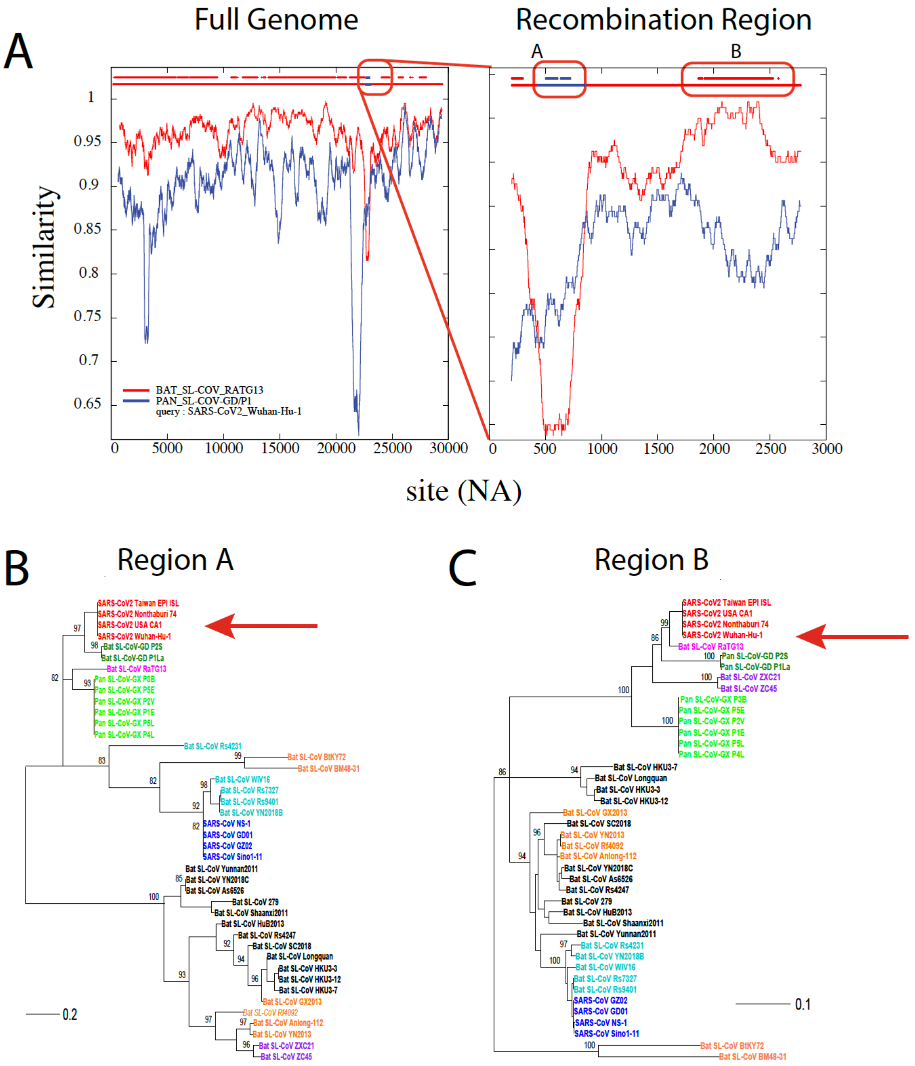


**Fig. S2. Recombination analysis of the CoV-SARS-2 Wuhan-Hu-1 sequence**. (A) Similarity plots comparing the Wuhan-Hu-1 sequence to the bat-Cov RaTG13 sequence (red) and the Pan_SL-CoV_GD/1PL sequence (blue). Plots were obtained using the LANL tool RIP using a window size of 400 bp. The full genome comparison is shown on the right and, on the left, a close-up of the recombination region is shown. Blue and red horizontal lines at the top of the panels show recombination breakpoints at the 99% confidence level. The blank between the A and B regions means uncertainty. (B and C) Phylogenetic trees of the individual recombination regions A and B, showing the different clusterings of the CoV-SARS-2 sequence compared to RaTG13 and Pan_SL-CoV_GD/1PL (highlighted by the red arrows).

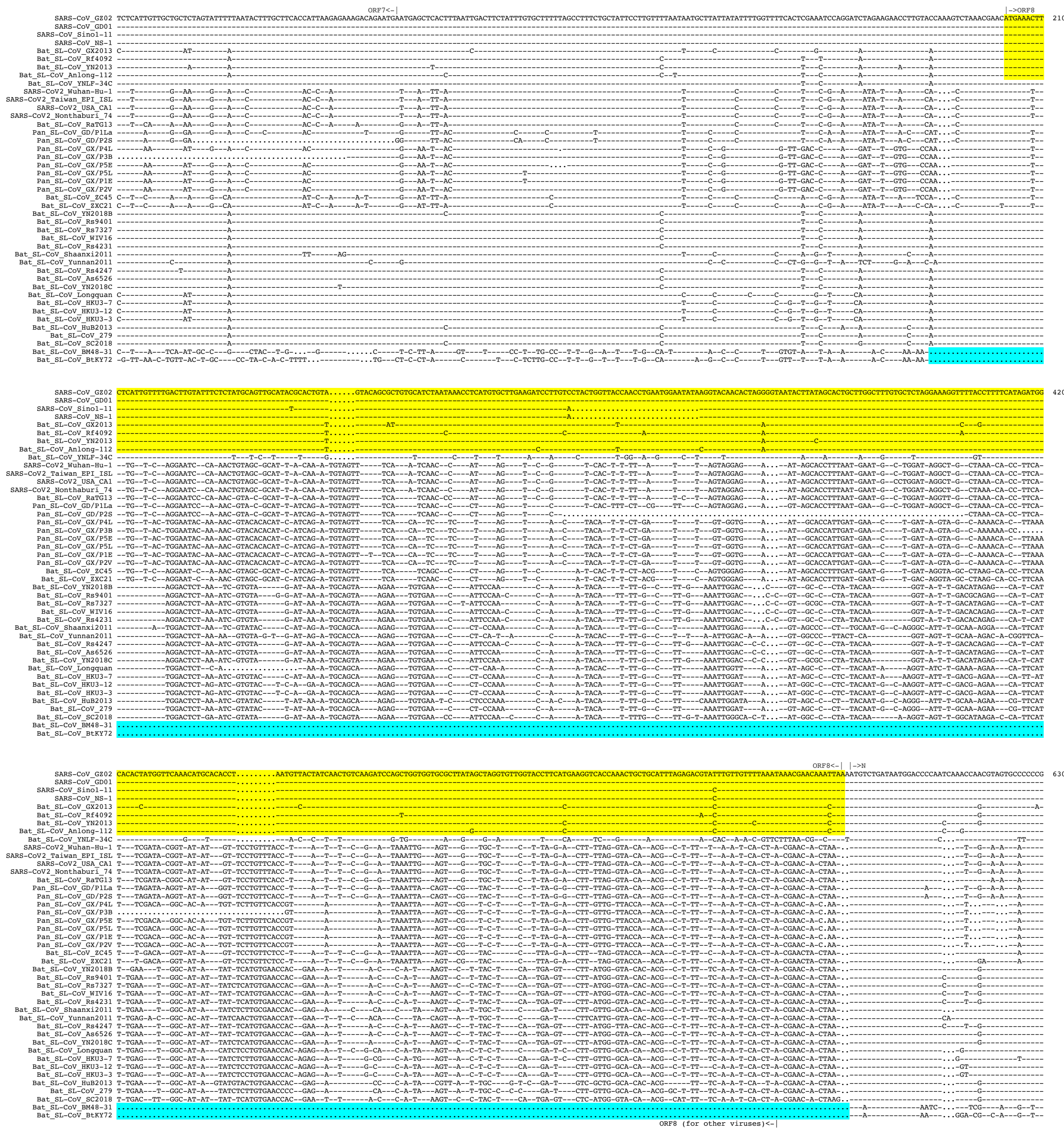

**Fig. S3. Highly conserved sequences around the receptor binding motif and furin cleavage sites among SARS-CoV-2, RaTG13 and Pan_SL-CoV viruses.** Alignment of amino acid sequences around receptor binding motif (RBM) and furin cleavage sites in the spike glycoprotein compared to Wuhan-Hu-1 (top sequence, na 22541-24391). Identical amino acids are shown as dashes and deletions as dots. RBM is shown at aa positions 439-508, and the furin cleavage site is highlighted in magenta. Regions with identical or nearly identical amino acid sequences among SARS-CoV-2, RaTG13 and Pan_SL-CoV viruses are highlighted in yellow. The positions of critical contact sites with ACE2 are indicated at the top of the alignment and highlighted in blue. The two large deletions in RBM are indicated in green.

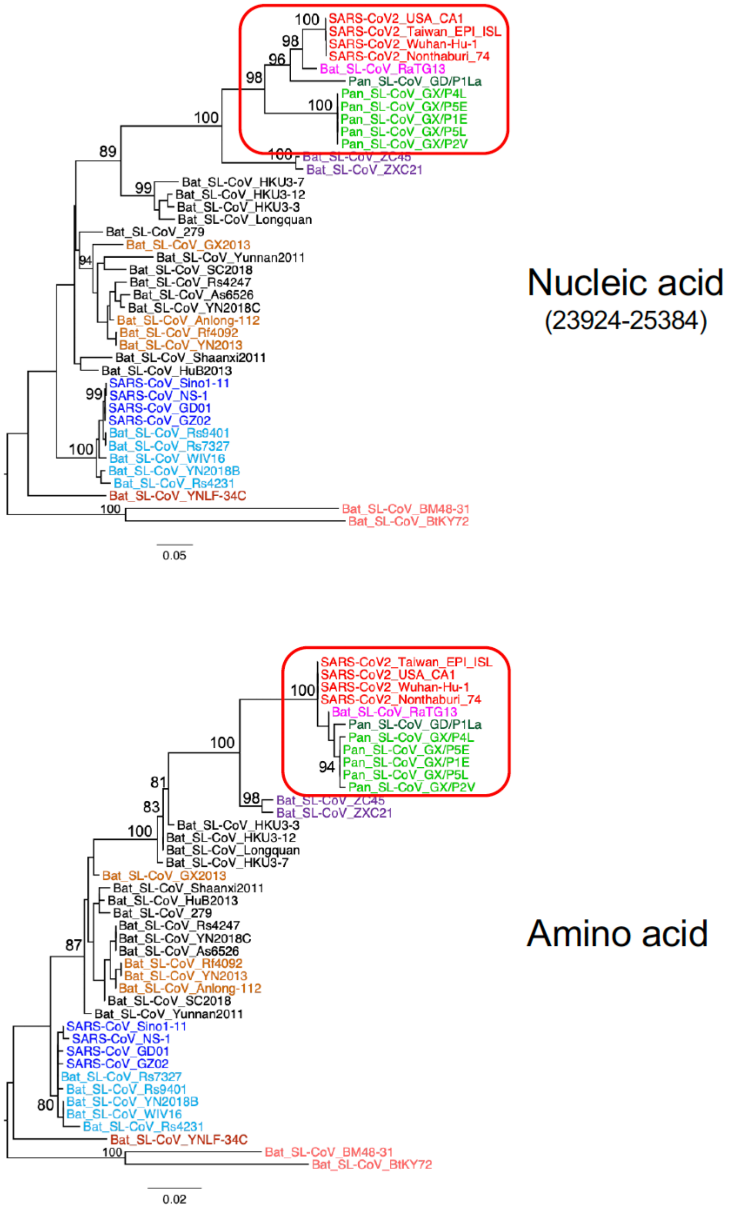


**Fig. S4. Purifying selection in the 3’ end region of the S gene.** Purifying selection pressure on the 3’ end region (na 23924-25384) of the S gene region among SARS-CoV-2, RaTG13 and Pan_SL-CoV viruses (within red boxes in phylogenetic trees) are shown by much shorter branches with amino acid sequences than with nucleic acid sequences.

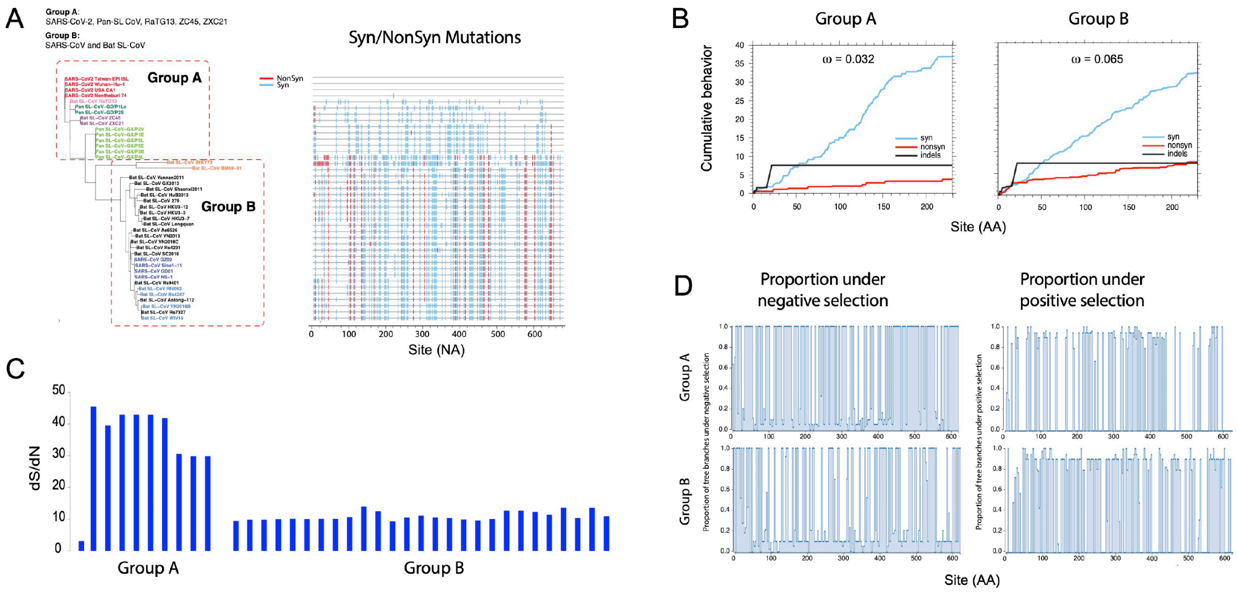


**Fig. S5. Purifying selection pressure on the M gene.** (A) Phylogenetic tree (left) and Highlighter plot (right) of all sequences compared to SARS-CoV-2 in the M gene. SARS-CoV-2, RaTG13, all Pan_SL-CoV and the two bat  CoV (ZXC21 and ZC45) sequences are in Group A, and all other sequences in Group B, to highlight differences between the two groups. Colored tic marks are mutations compared to the top sequence (SARS-CoV-2 Wuahn-Hu-1), with synonymous as light blue and non-synonymous as red. (B) Cumulative plots of the average behavior of each codon for all pairwise comparisons in the input data, for synonymous mutations, non-synonymous mutations and indels of group A sequences (left) and group B sequences (right). Average ratios of the rate of nonsynonymous substitutions per nonsynonymous site (dN/dS, or ω) for each sequence group are reported at the top of each plot. (C) dS/dN ratios of all sequences compared to Wuhan-Hu-1. (D) Proportion of tree branches under positive and negative selection (right and left respectively) for the two sequence groups as calculated using the mixed effects model of evolution (MEME) from the datamonkey (www.datamonkey.org) server.

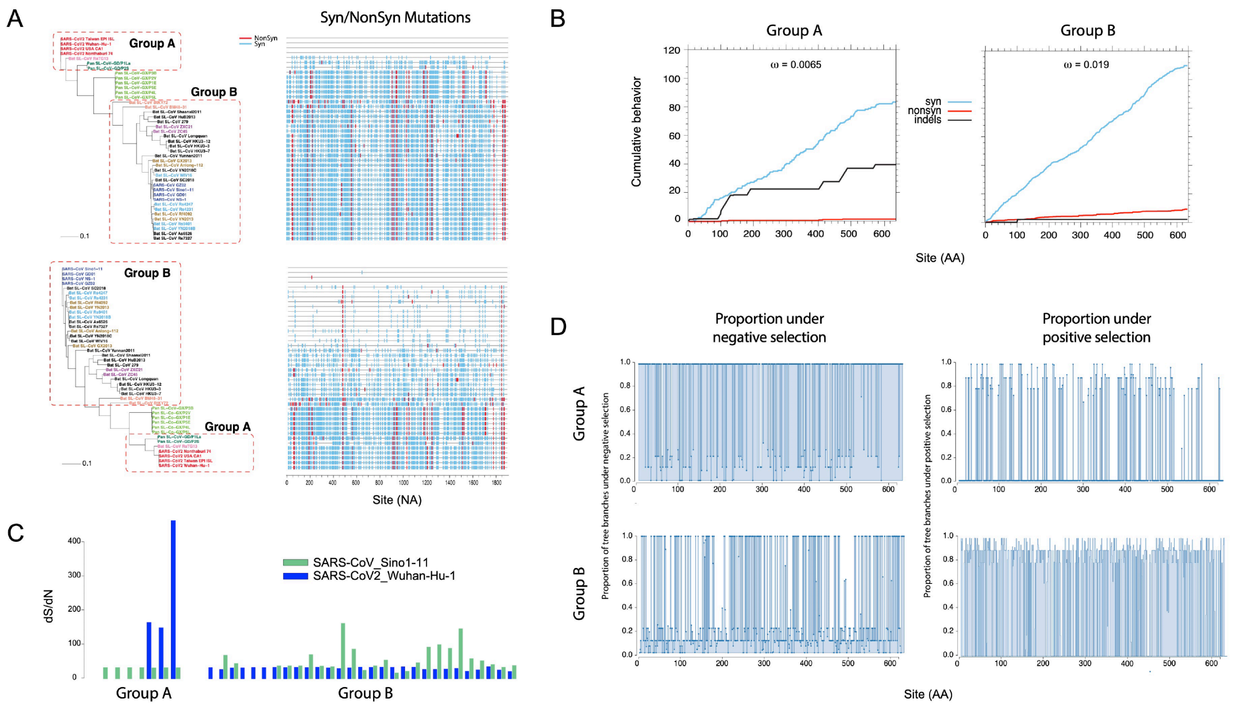


**Fig. S6. Purifying selection pressure on the partial region of ORF1a.** (A) Phylogenetic trees (left) and Highlighter plots (right) of sequences compared to SARS-CoV-2 (top) and to SARS-CoV (bottom) in the partial region of ORF1a. SARS-CoV-2, RaTG13 and Pan_SL-CoV from Guangdon are in Group A, and all other bat-CoV sequences in Group B, to highlight differences between the two groups. Colored tic marks are mutations compared to the top sequence (SARS-CoV-2 Wuahn-Hu-1 in the top graph and SARS-CoV Sin 1-11 in the bottom graph), with synonymous as light blue and non-synonymous as red. (B) Cumulative plots of the average behavior of each codon for all pairwise comparisons in the input data, for synonymous mutations, non-synonymous mutations and indels of group A sequences (left) and group B sequences (right). Average ratios of the rate of nonsynonymous substitutions per nonsynonymous site (dN/dS, or ω) for each sequence group are reported at the top of each plot. (C) dS/dN ratios of all sequences compared to Wuhan-Hu-1 in dark blue, and compared to SARS-CoV Sin 1-11 in green. (D) Proportion of tree branches under positive and negative selection (right and left respectively) for the two groups as calculated using the mixed effects model of evolution (MEME) from the datamonkey server (www.datamonkey.org).

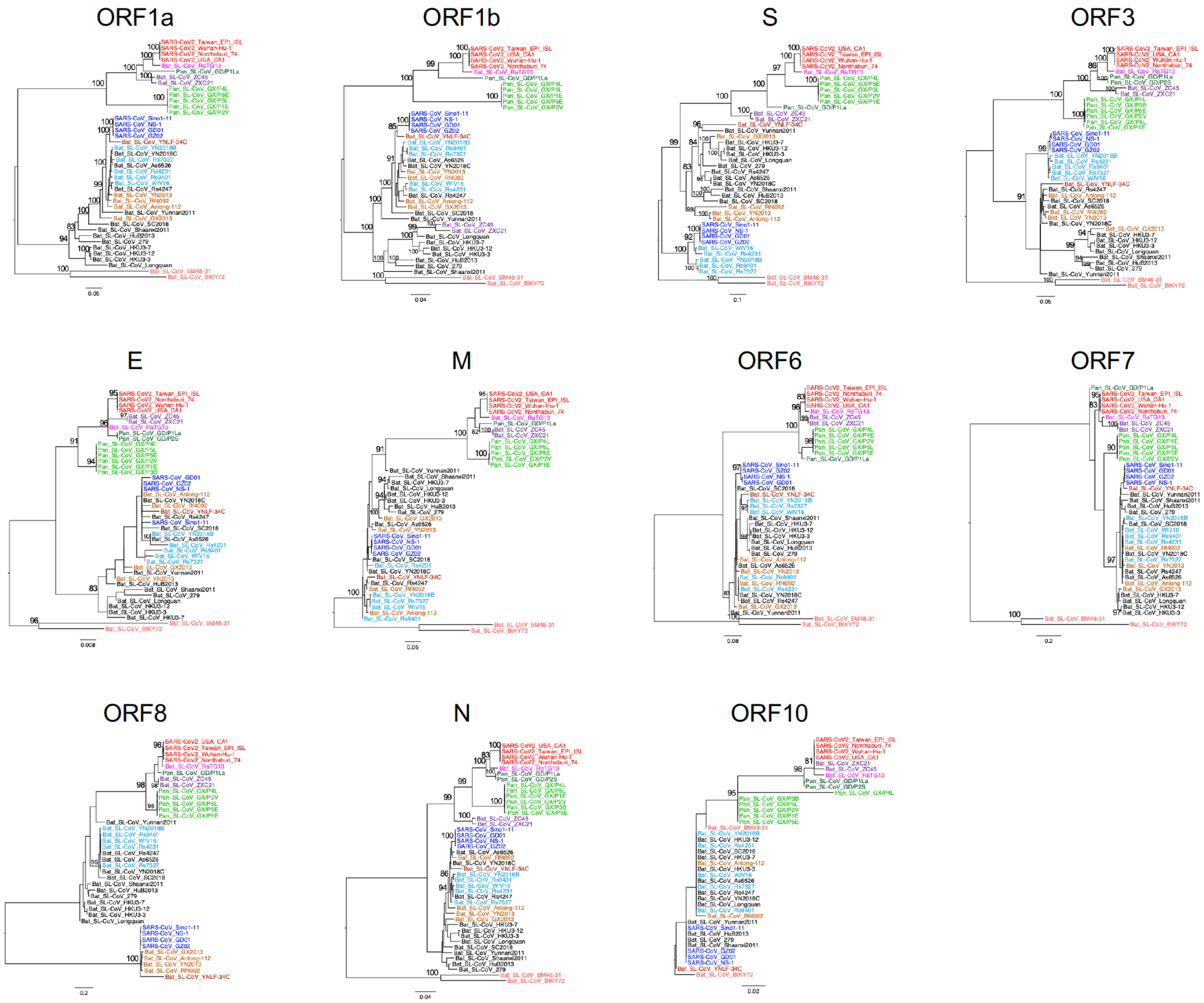


**Fig. S7. Phylogenetic tree analysis of SARS-CoV-2 genes together with other CoVs.** Phylogenetic trees were constructed for each coding region in the CoV genome. Sequences are colored differently based on their hosts and phylogenetic cluster: 4 SARS-CoV-2 sequences (red), Bat_SL-CoV sequence RaTG13 (magenta), 2 pangolin CoVs from Guangdong (Pan_SL-CoV_GD, dark green), 6 pangolin CoVs from Guangxi (Pan_SL-CoV_GX , light green), and 4 SARS-CoV sequences (dark blue). The remaining Bat_SL-CoV sequences in the set are color-coded according to their phylogenetic subclusterings  in the tree.

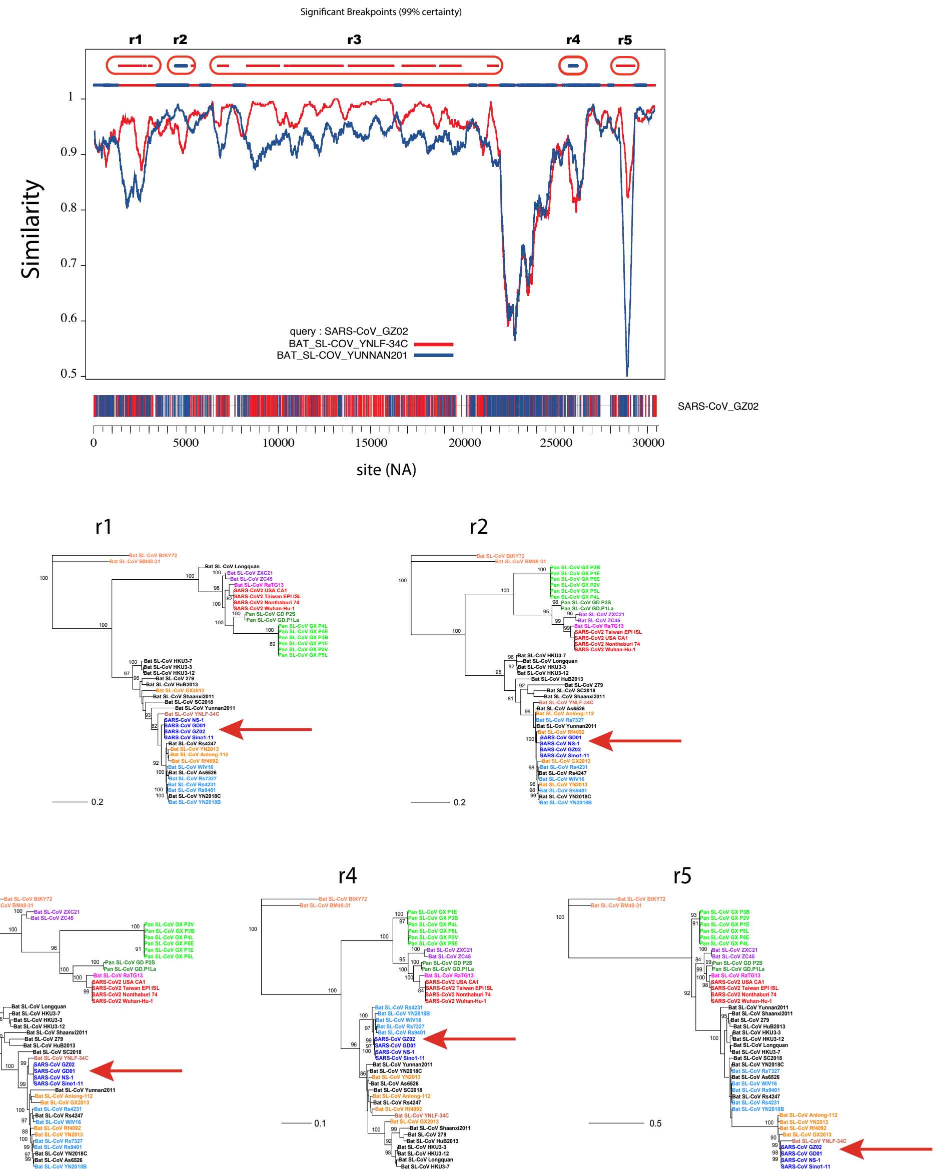


**Fig. S8. Recombination analysis of SARS-CoV sequences.** (A) Similarity plot comparing SARS sequence GZ02 to bat-CoV viruses YNLF-34C (red) and Yunnan2011 (blue). The plot was obtained using the recombination detection tool RIP with a window size of 400 base pairs (*10*). Top line shows break points at 99% confidence. Regions between significant break points are highlighted in the red ovals are marked r1-r5. At the bottom of the graph GZ02 is shown with nucleotide mutations colored in red if they are shared with sequence YNLF-34C, blue if they are shared with Yunnan2011. Nucleic acid unique to GZ02 are not shown. (B-F) Phylogenetic trees of the individual regions between break points, showing how the SARS sequences cluster more closely to either YNLF-34C or Yunnan2011 (red arrows). Regions between breakpoints were at the following base positions from the beginning of the genome: 1561-3303 (r1); 4621-5220 (r2); 5521-21360 (r3); 25201-25620 (r4); and 28201-29110 (r5).

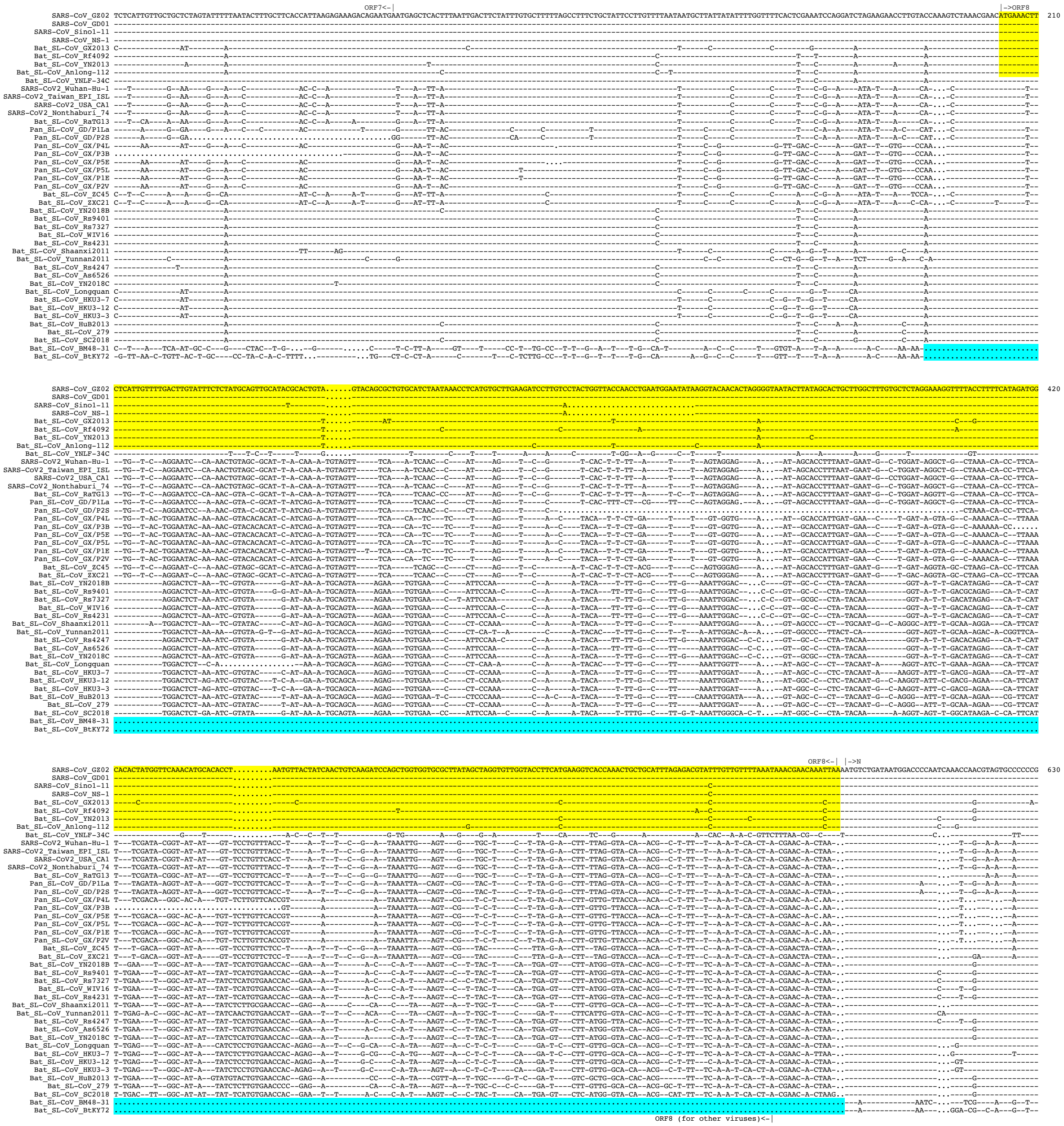

**Fig. S9. Nearly identical ORF8 sequences between SARS-CoV and four Bat_SL-CoV viruses.** Nucleic acid sequence alignments of ORF8, partial ORF7 and the N gene compared to SARS-CoV GZ02 (top). Identical amino acids are shown as dashes and deletions as dots. Regions with nearly identical sequences among SARS-CoV and four bat_SL-CoV (GX2013, Anlong-112, Rf4092 and YN2013) viruses are highlighted in yellow. The ORF8 deletion in two highly divergent bat-SL-CoVs (BtKY72 and BM48-31) is highlighted in blue.

**Table S1. Impact of amino acid substitutions in receptor binding motif**

| No. of mutation | Position in SARS2 RBM | AA in SARS2 | AA in RaTG13 | ∆∆G (kCal/Mol) | Effect for the RaTG13 mutations |
|---|---|---|---|---|---|
| 1 | | Asn | Lys | | No contact |
| 2 | | Asn | His | | No contact |
| 3 | | Leu | Ile | | No contact |
| 4 | | Ser | Ala | | No contact |
| 5 | | Val | Glu | | No contact |
| 6 | 449 | Tyr | Phe | 0.71 | Lost 1 h-bond |
| 7 | | Ser | Ala | | No contact |
| 8 | | Thr | Lys | | No contact |
| 9 | | Val | Gln | | No contact |
| 10 | | Glu | Thr | | No contact |
| 11 | 486 | Phe | Leu | 1.63 | Small/less hydrophobic |
| 12 | | Phe | Tyr | | No contact |
| 13 | 493 | Gln | Tyr | 3.44 | Gain a h-bond/too bulky |
| 14 | | Ser | Arg | | No contact |
| 15 | 498 | Gln | Tyr | 1.26 | too bulky |
| 16 | 501 | Asn | Asp | 0.31 | Buried a charge |
| 17 | | His | Tyr | | No contact |
